# *xbx-4*, a homolog of the Joubert syndrome gene FAM149B1, acts via the CCRK and MAK kinase cascade to regulate cilia morphology

**DOI:** 10.1101/2021.05.09.443182

**Authors:** Ashish K. Maurya, Piali Sengupta

## Abstract

Primary cilia are microtubule (MT)-based organelles that mediate sensory functions in multiple cell types. Disruption of cilia structure or function leads to a diverse collection of diseases termed ciliopathies (1–3). Mutations in the DUF3719 domain-containing protein FAM149B1 have recently been shown to elongate cilia via unknown mechanisms and result in the ciliopathy Joubert syndrome (4). The highly conserved CCRK and MAK/RCK kinases negatively regulate cilia length and structure in *Chlamydomonas, C. elegans*, and mammalian cells (5–11). How the activity of this kinase cascade is tuned to precisely regulate cilia architecture is unclear. Here we identify XBX-4, a DUF3719 domain-containing protein related to human FAM149B1, as a novel regulator of the DYF-18 CCRK and DYF-5 MAK kinase pathway in *C. elegans.* As in *dyf-18* and *dyf-5* mutants (11), sensory neuron cilia are elongated in *xbx-4* mutants and exhibit altered axonemal MT stability. XBX-4 promotes DYF-18 CCRK activity to regulate DYF-5 MAK function and localization. We find that Joubert syndrome-associated mutations in the XBX-4 DUF3719 domain also elongate cilia in *C. elegans.* Our results identify a new metazoan-specific regulator of this highly conserved kinase pathway, and suggest that FAM149B1 may similarly act via the CCRK/MAK kinase pathway to regulate ciliary homeostasis in humans.

## RESULTS and DISCUSSION

### XBX-4 acts in the DYF-5 kinase pathway to regulate cilia morphology

The integrity of cilia present on sensory neurons in the head amphid and tail phasmid organs in *C. elegans* can be readily assessed by the ability of a subset of these neurons to take up lipophilic dye (12, 13). Overexpression of DYF-5 MAK [*dyf-5(XS)*] severely truncates cilia resulting in a failure of neurons to dye-fill (5) (Figure S1). To identify regulators and effectors of DYF-5 activity, we performed a forward genetic screen and identified mutants that restored dye-filling in *dyf-5(XS)* animals (see Methods) (Figure 1A). In order to assess whether suppression of the dye-filling defect was correlated with restoration of cilia length in different neuron types in isolated mutants, we examined the cilia morphology of the non dye-filling AWA olfactory neurons. Wild-type AWA cilia exhibit complex and highly arborized morphologies (14, 15) (Figure 1B); these cilia are markedly truncated and lack all branches in *dyf-5(XS)* animals (11) (Figure 1B-C). We isolated twenty mutants in which both the dye-filling and truncated AWA cilia morphology phenotypes of *dyf-5(XS)* animals were significantly suppressed (Figure 1A-C, Figure S1).

**Figure 1.**
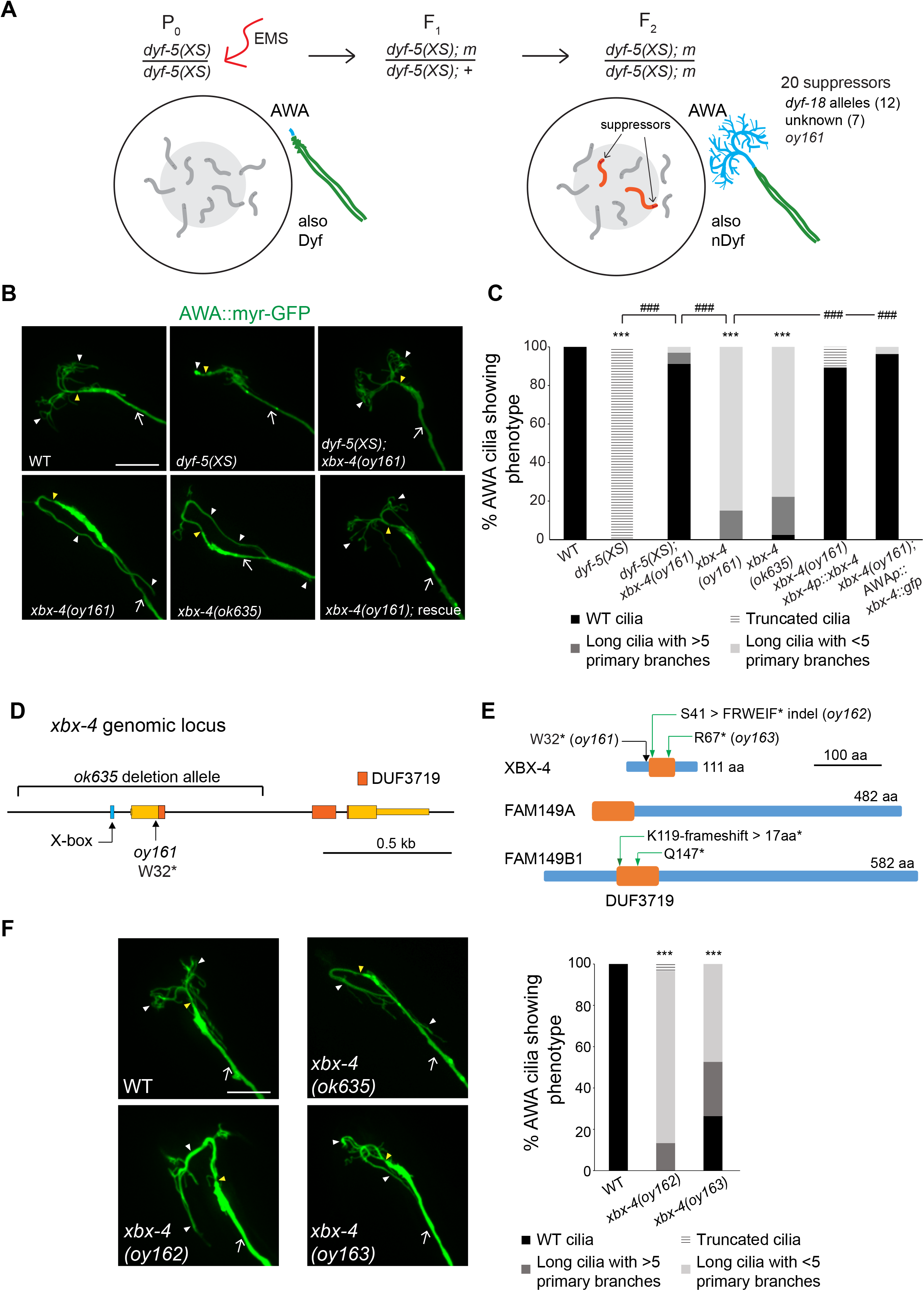
The XBX-4 DUF3719 containing protein regulates sensory cilia morphology and length. **A)**Schematic of forward genetic screen to identify *dyf-5(XS)* suppressors (5). Dyf: dye-filling defective, nDyf: non dye-filling defective; m: mutant; EMS: ethyl methanesulfonate. Animals carrying suppressor mutations were identified based on restoration of dye-filling and AWA cilia morphology. Also see Figure S1. **B-C)**Representative images (left) and quantification of AWA cilia phenotypes (right) in animals of the indicated genotypes. Rescue **(B)** indicates rescue with an *xbx-4*p*::xbx-4* transgene. The *gpa-4Δ6* promoter was used for AWA-specific rescue **(C)**. AWA was visualized using *gpa-4Δ6*p*::myr-gfp*. Anterior at top left. ***: different from wild-type at *P*<0.001; ###: different between indicated values at *P*<0.001 (Kruskal-Wallis test with Bonferroni post hoc corrections for multiple comparisons). n≥30 neurons each. Also see Figure S2. **D)**Structure of the *xbx-4* genomic locus. The locations of the *ok635* deletion and *oy161* nonsense alleles, and the X-box, are indicated. Sequences encoding the DUF3719 domain are shown. Thin yellow bar indicates untranslated sequences. **E)**Domains of XBX-4 and the human homologs FAM1491 and FAM149B1. Green arrows: locations of residues in FAM149B1 mutated in Joubert syndrome (4), and corresponding mutations engineered in the XBX-4 DUF3719 domain to generate *xbx-4(oy162)* and *xbx-4(oy163)* alleles. Also see Figure S3. **F)**Representative images (left) and quantification of AWA cilia morphologies in the indicated genetic backgrounds. ***: different from wild-type at *P*<0.001 (Kruskal-Wallis test with Bonferroni post hoc corrections for multiple comparisons). n≥30 neurons each. In all images, arrows: dendrite, yellow and white arrowheads: cilia base and cilia, respectively. Scale bar: 10 μm.

Complementation and targeted Sanger sequencing showed that twelve of these mutations were loss-of-function mutations in the *dyf-18* CCRK locus (Figure 1A). We previously showed that loss of *dyf-18* restores AWA cilia morphology and dye-filling in *dyf-5(XS)* animals (11), indicating that DYF-18 is necessary for maximal DYF-5 activity. Seven additional mutants exhibited no dye-filling or AWA cilia morphology defects in the absence of the *dyf-5(XS)* transgene. These mutations may affect genes regulating *dyf-5* promoter activity or *dyf-5(XS)* transgene expression and were not followed further (Figure 1A). The remaining *oy161* mutation complemented *dyf-18(ok200)* for both suppression of dye-filling and AWA cilia morphology phenotypes (data not shown). While *oy161* mutants alone in the absence of the *dyf-5(XS)* transgene exhibited only weak dye-filling defects (Figure S1), these mutants exhibited severely elongated and largely unbranched AWA cilia, similar to the phenotypes of *dyf-5* and *dyf-18* loss-of-function mutants (Figure 1B-C) (11).

Single nucleotide polymorphism mapping and whole genome sequencing (16) identified a nonsense mutation in the *xbx-4* (X-box promoter element regulated) gene in the *oy161* strain (Figure 1D) (17). Similar to *oy161* mutants, animals carrying the independently isolated *xbx-4(ok635)* deletion mutation (17) (Figure 1D) also exhibited elongated and unbranched AWA cilia, weak dye-filling defects, and failed to complement *oy161* for this phenotype (Figure 1B-C, Figure S1, and data not shown). The AWA cilia phenotype of *oy161* mutants was robustly rescued upon expression of wild-type *xbx-4* genomic sequences (Figure 1B-C). Moreover, expression of *xbx-4* sequences under the AWA-specific *gpa-4Δ6* promoter was also sufficient to rescue the *oy161* mutant phenotype in AWA (Figure 1C). We conclude that *oy161* is a mutation in the *xbx-4* gene, and that this gene acts in the DYF-5 kinase pathway to regulate cilia morphology and length.

In addition to AWA, the cilia of the AWB and AWC amphid sensory neurons exhibit complex morphologies, whereas additional amphid sensory neurons present within a channel formed by glia contain simpler rod-like cilia (channel cilia) (14, 15). While channel cilia are slightly, albeit significantly, elongated in *dyf-18* and *dyf-5* mutants (5, 11, 18), complex cilia morphologies are more dramatically affected in these mutant backgrounds (11). *xbx-4(ok635)* mutants also exhibited elongated AWB and AWC cilia (Figure S2A), whereas the rod-like cilia of the ASH neurons exhibited more minor, but statistically significant, elongation (Figure S2B). Together, these results indicate that XBX-4 regulates cilia morphology and length in multiple amphid sensory neuron types.

### *xbx-4* encodes a DUF3719 domain-containing protein related to the Joubert syndrome-associated protein FAM149B1

*xbx-4* was previously identified in an *in silico* screen for genes containing the predicted binding site (X-box) for the DAF-19 RFX transcription factor required for ciliogenesis (17, 19). Although *xbx-4* expression was shown to be regulated by DAF-19, no obvious ciliary or sensory abnormalities were previously reported in *xbx-4* mutant animals (17). *xbx-4* encodes a novel small protein of 111 aa containing a conserved metazoan-specific 70 aa domain of unknown function 3719 (DUF3719) (Figure 1D). The *oy161* mutation truncates the protein just N-terminal to the predicted DUF3719 domain, whereas the *ok635* mutation deletes upstream regulatory sequences including the X-box and part of the DUF3719 domain (Figure 1D, Figure S3A). The DUF3719 domain is present in only two proteins predicted to be encoded by the human genome – FAM149A and FAM149B1 (Figure 1E, Figure S3A). Sequence analysis indicated that the DUF3719 domain of XBX-4 is more closely related to the domain present in FAM149B1 (Figure S3B). However, both human proteins are larger than XBX-4 and contain additional residues outside the DUF3719 domain that are not shared with XBX-4 (Figure 1E, Figure S3A). While the function of FAM149A is unknown, mutations in the DUF3719 domain of FAM149B1 (Figure 1E, Figure S3A) have recently been shown to underlie phenotypes associated with the ciliopathy Joubert syndrome (4). Patient fibroblasts exhibit longer cilia and altered ciliary signaling (4), although the pathway by which FAM149B1 acts to regulate cilia length and function is unknown. These observations suggest a conserved role for the DUF3719 domain in regulating cilia architecture.

Joubert syndrome-associated mutations in FAM149B1 include a 2bp deletion that results in an early frameshift and truncates ~90% of the DUF3719 domain (K119-frameshift > 17aa > STOP), and a nonsense mutation that truncates ~50% of this domain (Q147 > STOP) (Figure 1E, Figure S3A). Since the DUF3719 domain is present at the N-terminus of FAM149B1 (Figure 1E), these mutations also delete C-terminal residues of this protein that are not conserved in XBX-4. We asked whether similar truncations of the DUF3719 domain in XBX-4 also result in ciliary defects in *C. elegans*. Via gene editing at the endogenous *xbx-4* locus, we generated the *xbx-4(oy162)* allele that is predicted to encode a protein in which the S41 residue is mutated to mimic the early frameshift/truncation mutation in human FAM149B1 (Figure 1E, Figure S3A). We also mutated R67 in XBX-4 (corresponding to Q147 in FAM149B1) to a STOP (*oy163*), resulting in loss of ~50% of the XBX-4 DUF3719 domain (Figure 1E, Figure S3A). *xbx-4(oy162)* animals exhibited strong AWA ciliary morphology defects resembling those in *xbx-4(ok635)* null mutants, whereas *xbx-4(oy163)* mutants exhibited similar but weaker phenotypes (Figure 1F). We infer that the DUF3719 domain is critical for XBX-4, and likely FAM149B1, function.

### XBX-4 regulates axonemal MT stability

The mechanisms by which FAM149B1 regulates cilia length in humans are unknown. The possible related functions of XBX-4 and FAM149B1 provide an opportunity to use the *C. elegans* system to characterize the pathways in which these proteins act to regulate cilia structure and function. We previously showed that the elongated AWA cilia phenotype in *dyf-18* and *dyf-5* mutants arises in part due to increased axonemal MT stability in these mutant backgrounds (11). While the end-binding protein EBP-2 is localized to the primary proximal stalk and distal tips of the branches in wild-type AWA cilia, this protein is present along the entire elongated AWA cilium and not enriched at the distal ciliary endings in *dyf-18* and *dyf-5* mutants (11). We also showed that destabilizing temperature-sensitive missense mutations in the *tbb-4* and *tba-5* tubulin genes (20) suppress the elongated cilia and loss of branching phenotype of *dyf-18* mutants at 15°C (11). Given the similar ciliary phenotypes of *xbx-4, dyf-5* and *dyf-18* mutants, we tested whether AWA ciliary axonemal MTs are similarly affected in *xbx-4* mutants.

As in *dyf-5* and *dyf-18* mutants (11), both EBP-2::GFP and TBB-4::tagRFP were localized along the entire AWA axoneme in *xbx-4(ok635)* mutants (Figure 2A). Moreover, the *tbb-4(sa127)* missense mutation significantly suppressed the elongated cilia phenotype of *xbx-4(ok635)* mutants such that a large majority of *xbx-4(ok635); tbb-4(sa127)* mutants exhibited wildtype-like branched AWA cilia (Figure 2B). As in *dyf-18* mutants but distinct from the localization patterns in wild-type animals (11), the ciliary intraflagellar transport B (IFT-B) protein OSM-6::GFP was localized at low levels in the elongated AWA cilium in *xbx-4* mutants (Figure 2C). We conclude that similar to our previous observations in *dyf-18* and *dyf-5* mutants, the elongated AWA cilia phenotype in *xbx-4* mutants arises in part due to increased axonemal MT stability.

**Figure 2.**
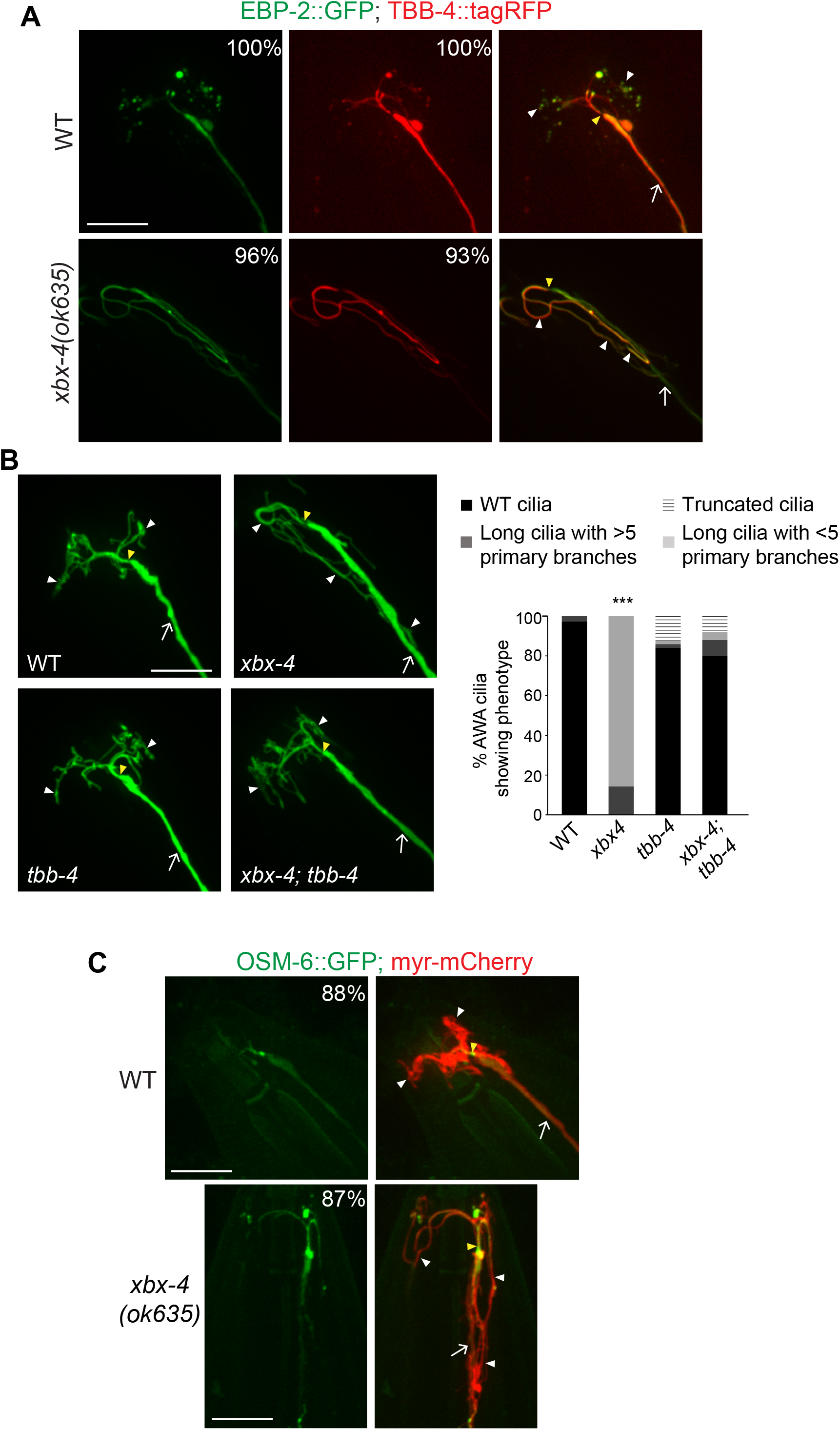
Axonemal microtubule dynamics may be altered in *xbx-4* mutants. **A,C)**Representative images of AWA cilia in the indicated genetic backgrounds. Numbers at top right indicate the percentage of neurons showing the phenotype. AWA was visualized using *gpa-4Δ6*p*::myr-mCherry* **(C)**. EBP-2::GFP, TBB-4::tagRFP **(A)** and OSM-6::GFP **(C)** were expressed under *gpa-4Δ6* regulatory sequences. n≥30 neurons each. **B)**Representative images (left) and quantification (right) of AWA cilia morphology in the shown genetic backgrounds. Alleles used were *xbx-4(ok635)* and *tbb-4(sa127).* Animals were grown at 15°C. AWA was visualized using *gpa-4Δ6*p*::myr-gfp.* ***: different from wild-type at *P*<0.001 (Kruskal-Wallis test with Bonferroni post hoc corrections for multiple comparisons). n≥30 neurons each. In all images, arrows: dendrite, yellow and white arrowheads: cilia base and cilia, respectively. Anterior at top left. Scale bar: 10 μm.

### XBX-4 acts upstream of DYF-18

The similar ciliary phenotypes of *xbx-4, dyf-18,* and *dyf-5* loss of function mutants, together with the observation that mutations in either *dyf-18* or *xbx-4* suppress the AWA ciliary phenotype of *dyf-5(XS)* animals to a comparable extent (Figure 1B-C) (11), suggest that XBX-4 acts in, or in parallel to, the DYF-18 and DYF-5 kinase pathway to regulate AWA cilia morphology.

We first tested whether loss of *xbx-4* enhances the AWA cilia phenotype of *dyf-5* or *dyf-18* single mutants. However, the AWA cilia morphology defects of *xbx-4 dyf-18* or *dyf-5; xbx-4* double mutants were no more severe than those of *dyf-18* or *dyf-5* single mutants alone (Figure 3A). Moreover, the AWA cilia morphology defects of *dyf-5; xbx-4 dyf-18* triple mutants were also not significantly different from those of *dyf-5; dyf-18* double mutants (Figure 3A). Although these observations suggest that XBX-4 acts in a linear pathway with DYF-18 and DYF-5, it remains possible that AWA cilia length was not further enhanced in the double and triple mutants due to a ceiling effect.

**Figure 3.**
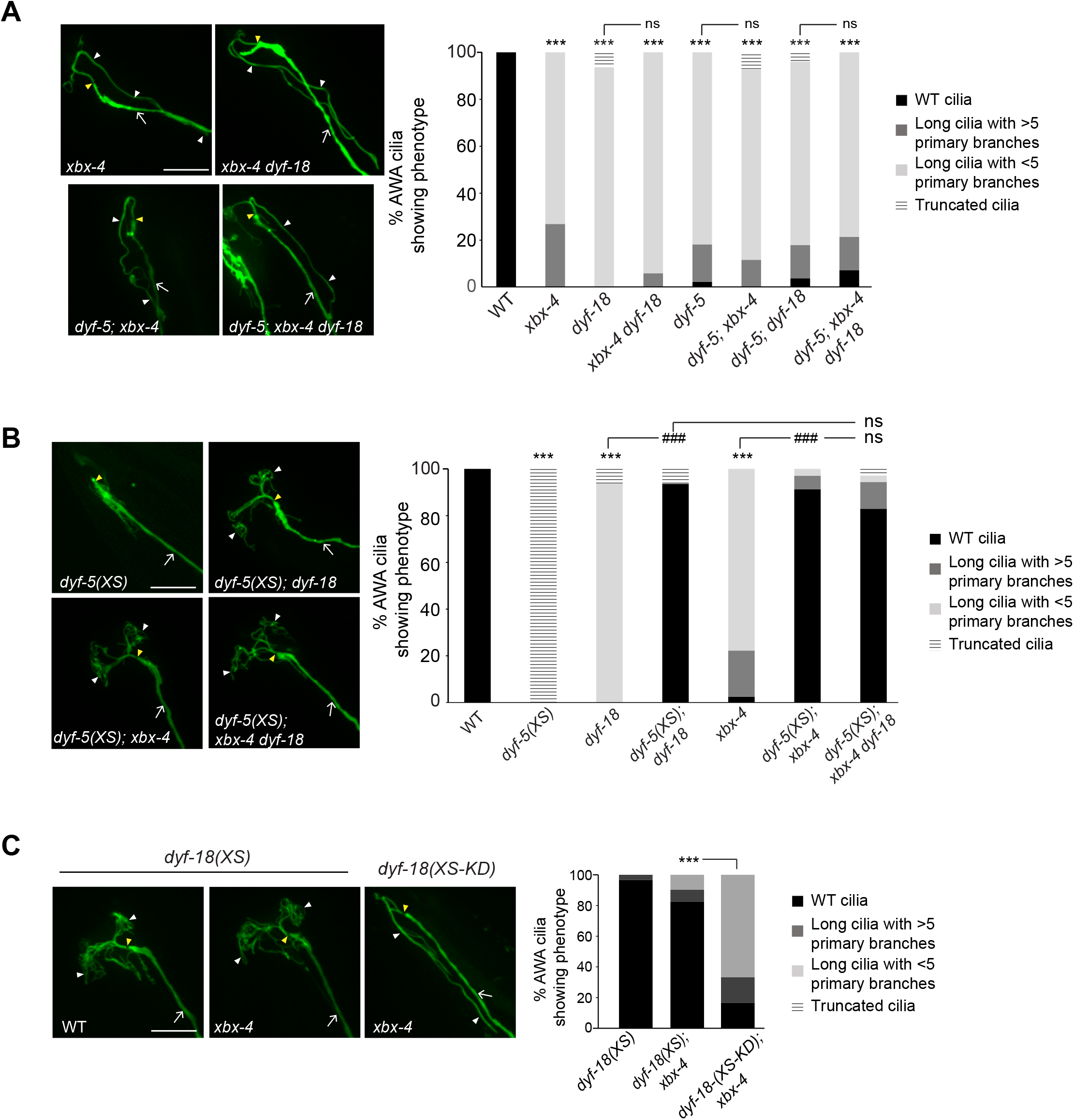
XBX-4 acts upstream of DYF-18 to regulate DYF-5. **A-C)**Representative images (left) and quantification (right) of AWA cilia morphology in the indicated genetic backgrounds. AWA was visualized using *gpa-4Δ6*p*::myr-gfp*. Alleles used were *dyf-18(ok200)* and *xbx-4(ok635)*. KD: kinase-dead. ***: different from wild-type at *P*<0.001, ###: different from indicated at *P*<0.001; ns: not significant (Kruskal-Wallis test with Bonferroni post hoc corrections for multiple comparisons). n≥30 neurons each. In all images, arrows: dendrite, yellow and white arrowheads: cilia base and cilia, respectively. Anterior at top left. Scale bar: 10 μm.

To address this issue, we next examined AWA cilia in *dyf-5(XS)* animals mutant for both *xbx-4* and *dyf-18*. We previously proposed that DYF-18 is necessary to maximally activate overexpressed DYF-5 and promote cilia truncation (11). In *dyf-18* mutants, DYF-5(XS) activity is reduced but not abolished, such that the AWA cilia morphology of *dyf-5(XS); dyf-18* double mutants is similar to that in wild-type animals (11). If XBX-4 acts in parallel to DYF-18 to regulate DYF-5, loss of both *xbx-4* and *dyf-18* together should further reduce DYF-5(XS) activity possibly phenocopying *dyf-5(lof)* mutants. However, we found that AWA cilia phenotype of the *dyf-5(XS); xbx-4 dyf-18* triple mutants was indistinguishable from that of *dyf-5(XS); dyf-18(ok200)* double mutants (Figure 3B). These results suggest that XBX-4 likely acts upstream or downstream of DYF-18 to regulate DYF-5 activity in AWA. Since CCRK directly phosphorylates MAK *in vitro* and in HEK293 cells (21, 22), we suggest that XBX-4 acts upstream of DYF-18.

To further test this hypothesis, we asked whether overexpression of DYF-18 is sufficient to suppress the AWA ciliary phenotype of *xbx-4(ok635)* mutants. Overexpression of *dyf-18* in ciliated neurons via the *bbs-8* promoter resulted in significantly shorter and more branched AWA cilia in *xbx-4* mutants, closely resembling the morphology of wild-type cilia (Figure 3C). In contrast, overexpression of a kinase-dead DYF-18 had little or no effect on the AWA cilia phenotype of *xbx-4* mutants (Figure 3C). These results further support the notion that XBX-4 functions upstream of, and regulates, DYF-18 function.

### XBX-4 acts via DYF-18 to regulate DYF-5 localization

To begin to investigate how XBX-4 might modulate DYF-18, we first examined the expression pattern and subcellular localization of XBX-4. An *xbx-4* cDNA tagged with GFP-encoding sequences at the C-terminus and driven under *xbx-4* upstream regulatory sequences was expressed in the majority of ciliated sensory neurons in the head (Figure 4A) (17). An XBX-4::GFP fusion protein expressed specifically in AWA showed diffuse localization throughout the soma and processes (Figure 4A) and robustly rescued the AWA cilia phenotype of *xbx-4* mutants (Figure 1C). In AWA, XBX-4::GFP was present throughout the neuron including in the main proximal ciliary stalk with weaker expression in the distal ciliary branches (Figure 4A). This localization pattern resembled that of DYF-18::GFP in AWA, but was distinct from that of GFP::DYF-5 which is primarily enriched in the proximal stalks of AWA cilia (11). A rescuing GFP::XBX-4 protein did not undergo IFT (Figure S4A-B), and localization of this protein appeared unaltered in *dyf-18* or *dyf-5* mutants (Figure S4C).

**Figure 4.**
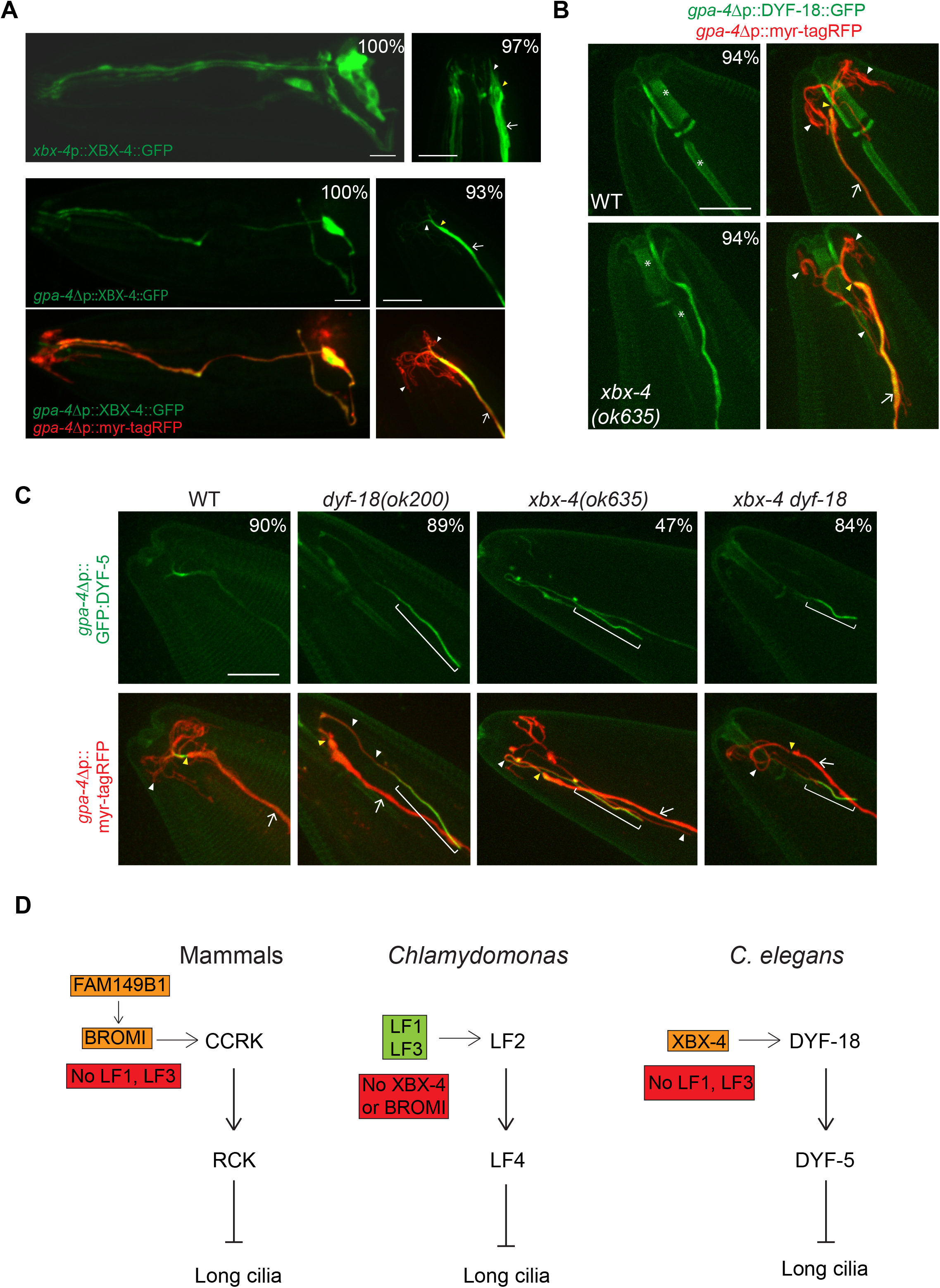
XBX-4 regulates DYF-5 localization in AWA. **A)**Representative images of XBX-4 localization in ciliated sensory neurons. XBX-4::GFP was expressed under its own promoter or under AWA-specific *gpa-4Δ6* regulatory sequences. Images show expression in the entire neuron(s) (left) and in the distal dendrites and cilia (right). Images at left and right are from different animals. Numbers at top right indicate percentage of neurons exhibiting the phenotype. AWA was visualized using *gpa-4Δ6*p*::myr-tagRFP*. Anterior at left (left) or at top (right). n≥30 neurons each. **B)**Representative images of DYF-18::GFP localization in AWA in WT and *xbx-4* mutants. Expression in AWA was driven under the *gpa-4Δ6* promoter. AWA was visualized using *gpa-4Δ6*p*::myr-tagRFP*. Asterisks indicate autofluorescence in the pharynx. Numbers at top right indicate the percentage of neurons showing the phenotype. Anterior at top left. n≥30 neurons each. **C)**Localization of GFP::DYF-5 expressed in AWA in the indicated genetic backgrounds. Brackets indicate the distal tip of the AWA cilium. Numbers at top right indicate the percentage of neurons showing the phenotype. Anterior at top left. n≥30 neurons each. In all images, arrows: dendrite, yellow and white arrowheads: cilia base and cilia, respectively. Scale bar: 10 μm. **D)**Schematic of known regulators and interactors of the CCRK and MAK kinase pathway in different organisms. See text for details and references.

We next tested the hypothesis that XBX-4 regulates DYF-18 localization and/or activity. Loss of *xbx-4* had no effect on the level or localization of DYF-18::GFP expressed specifically in AWA (Figure 4B). DYF-18 activity can be assessed by examining localization of the downstream DYF-5 kinase. While GFP::DYF-5 localization is restricted to the proximal stalk of wild-type AWA cilia, this fusion protein is instead present throughout the elongated and unbranched AWA cilia with enrichment at the distal tip in *dyf-18* mutants (Figure 4C) (11). We found that the localization pattern of GFP::DYF-5 in AWA cilia in *xbx-4* mutants was similar to that in *dyf-18* mutants (Figure 4C). Loss of both *xbx-4* and *dyf-18* together did not further alter GFP::DYF-5 localization (Figure 4C). We infer that XBX-4 regulates DYF-18 activity to control DYF-5 localization and AWA cilia architecture.

In summary, our results identify XBX-4 as a new metazoan-specific regulator of the highly conserved DYF-18 CCRK and DYF-5 MAK kinase cascade implicated in the regulation of cilia length and morphology across species (Figure 4D). Given the related sequences and phenotypic consequences of mutations in the DUF3719 domains of XBX-4 and FAM149B1, our observations raise the possibility that FAM149B1 also acts via the CCRK/MAK pathway to maintain cilia length in humans. Despite the functional conservation of the core CCRK and MAK regulatory pathway in ciliary length control, this pathway appears to interact with, and be regulated by, diverse molecules in different species. For instance, the LF1 and LF3 proteins interact with LF2 CCRK in *Chlamydomonas,* and *lf1* and *lf3* mutants also exhibit elongated flagella (8, 23, 24). However, LF1 and LF3 homologs are not present in mammals or *C. elegans*, and conversely, the *Chlamydomonas* genome does not encode proteins containing the DUF3719 domain. The FAP20 protein was identified as a CCRK interactor in human interactome studies (25, 26), but this protein appears to have distinct ciliary roles in different species and cilia types (27–29). CCRK also directly interacts with the Broad-minded (BROMI) protein in vertebrates, and BROMI was shown to regulate CCRK stability (30, 31). BROMI in turn interacts with FAM149B1 (4). The *Chlamydomonas* genome does not encode a Bromi homolog, and mutations in a homolog in *C. elegans* do not affect cilia (A.K.M. and P.S., unpublished). The concept of interconnected functional modules in cellular networks is well-established (32, 33). We propose that the CCRK and MAK kinases together form a functional module whose output is tuned by diverse regulatory inputs to permit precise shaping of cilia morphologies in a species-, cell-, and context-dependent manner (Figure 4D).

A characteristic of ciliopathies such as Joubert Syndrome is their pleiotropy and variable expressivity across cell and tissue types (2, 34, 35). Specific mutations in ciliary genes affect different cells and tissues to different extents, highlighting the context-specific roles of these conserved proteins. Thus, while mutations in *dyf-5, dyf-18* and *xbx-4* result in relatively minor effects on cilia length in channel cilia in *C. elegans* possibly due to anatomical constraints (5, 11, 18) (this work), AWA and other complex wing cilia are dramatically elongated in these mutants (11) (this work). Moreover, while DYF-18 also regulates the localization of the CDKL-1 kinase in amphid channel cilia to influence cilia length (36), AWA cilia morphology is unaffected in *cdkl-1* mutants (A.K.M. and P.S., unpublished). *C. elegans* is now well-established as an excellent experimental system in which to explore the roles of genes implicated in ciliopathies (37–39). Examining the roles of ciliopathy-associated genes across diverse ciliated species and cell types will allow for a more complete understanding of the underlying pathways, and provide insights into how cell type-specific modulation of conserved ciliogenic modules drives structural and functional specialization of cilia in development and disease.

## METHODS

### Growth of *C. elegans*

*C. elegans* strains were grown on *E. coli* OP50 bacteria using standard procedures. The wild-type strain was *C. elegans* Bristol N2. Double mutant strains were generated using standard genetic methods, and the presence of the desired mutations was verified by PCR-based genotyping and/or sequencing. Animals were maintained with plentiful food for at least two generations at 20°C before analyses unless indicated otherwise. All strains used in this work are listed in Table S1.

### Generation of transgenic strains

DNA constructs were injected at 5-30 ng/μl. *unc-122*p::*gfp* or *unc-122*p::*mCherry* injected at 15-30 ng/μl were used as coinjection markers. All shown data are from two independent transgenic lines each unless indicated otherwise. Transgenes expressed from the same extrachromosomal array were examined in wild-type and mutant animals.

### Isolation of *dyf-5(XS)* suppressors

*dyf-5(XS)* suppressors were identified in a forward genetic screen. Animals from the strain PY11440 (Table S1) carrying stably integrated *dyf-5(XS)* and *gpa-4Δ6*p::*myr-gfp* transgenes were mutagenized with EMS using standard procedures, and 8-12 P0 animals were placed on each of 8 NGM growth plates. Subsequently, 4-6 F1 animals from each P0 plate were individually cloned out onto 10 NGM growth plates, and their F2 progeny examined for dye-filling as described below. Animals identified as defective in dye-filling were recovered from the imaging slide and their progeny were examined for their AWA cilia morphology phenotype. Twenty mutants were isolated in which both the dye-filling and AWA cilia morphology defects of *dyf-5(XS)* were significantly restored.

### Cloning of *xbx-4(oy161)*

The *oy161* allele identified in the forward genetic screen was outcrossed to N2 and segregated from the *dyf-5(XS)* transgene. The unbranched and long AWA cilia phenotype of *oy161* mutants was used to map and identify the affected gene. *oy161* was mapped to LG I using the EG8040 and EG8041 fluorescent mapping strains (40). To identify the causal mutation, *oy161* animals carrying the stably integrated *gpa-4Δ6*p::*myr-gfp* transgene were crossed to the polymorphic Hawaiian CB4856 strain. Around 60 F2 progeny of this cross exhibiting elongated and unbranched AWA cilia were identified (homozygous mutants) using a compound fluorescence microscope, genomic DNA was isolated from their pooled progeny, and subjected to Illumina (paired end 150 bp) sequencing (20X coverage, Beijing Genomics Institute) (16).

The density of unique and polymorphic single nucleotide polymorphisms was assessed using online tools (CloudMap) on usegalaxy.org server as previously described (41). The generated chromosomal level frequency plots (bins of 1 Mb and 0.5 Mb) for pure parental alleles showed a single peak of unique variants on chromosome 1. Within the middle of this peak a nonsense mutation in *xbx-4* was identified (*oy161*). An independently obtained deletion allele (*ok635*; CGC) displayed similar AWA ciliary phenotypes and failed to complement *oy161*.

### Dye-filling

Animals were washed from culture plates into 1.5 ml tubes, and pelleted and resuspended in 100-200 μl of M9 with 10 μg/ml DiI (Life Technologies). Tubes were gently rocked for 2 hrs and animals were subsequently washed with M9 and placed on seeded NGM plates. Dye-filling was assessed after 20 mins under a fluorescent dissecting microscope.

### Microscopy

Cilia were imaged by anesthetizing animals with 10 mM tetramisole hydrochloride (Sigma) and mounted on agarose pads (2-10%). Animals were imaged using a 100X objective on a spinning disk confocal microscope (Zeiss Axio Observer with a Yokogawa CSU-22 spinning disk confocal head). Confocal sections were obtained in 0.25 μm steps in the *z*-axis and maximum intensity projections were obtained using SlideBook 6.0 software (3i, Intelligent Imaging Innovations). To visualize cilia, images were adjusted in ImageJ (NIH) for brightness and contrast.

### Generation of *xbx-4(oy162)* and *xbx-4(oy163)* alleles

*xbx-4(oy162)* and *xbx-4(oy163)* were generated via CRISPR-Cas9-mediated gene editing as described (42). crRNA and homology oligo donors were obtained from Integrated DNA Technologies (IDT). The edited sequences were designed to include EcoRV and NheI restriction sites. Animals were injected with Cas9-crRNA RNP complexes along with the oligo homology donor and a *unc-122*p*::gfp* co-injection marker. Animals transgenic for the coinjected marker were isolated and their progeny analyzed for the presence of the desired mutations by sequencing. *xbx-4(oy162)* crRNA: 5’-<u>TTCGATGGGAAAAGTACTGG</u>GTTTTAGAGCTATGCT-3’ *xbx-4(oy162)* oligo donor: 5’-GTTTCTGGATTCATGGGATAACGCGGTTTTCGATGGGtAAAGTACTGGTGgtgggtggctttttttttgaaaatagtaat-3’ *xbx-4(oy163)* crRNA: 5’-<u>CTGAGATTCAAAACGGGATC</u>GTTTTAGAGCTATGCT-3’ *xbx-4(oy163)* oligo donor: 5’-TATGGAGGGACAAATTTCCTCATTTAAGGATTACCGGA<u>agCtagc</u>TTGAATCTCAGAAA GTGACGgtaataagaatttttact-3’

### Molecular biology

To generate *xbx-4*p*::xbx-4*, *xbx-4* coding sequences were amplified from genomic DNA and cloned into the pPD95.77 expression vector along with 1293 bp upstream and 944 bp downstream *xbx-4* genomic sequences using Gibson assembly (NEBuilder® HiFi DNA Assembly Master Mix). N- and C-terminal GFP-tagged XBX-4 was generated by inserting GFP coding sequences at the appropriate location using PCR and Gibson cloning. *gfp::xbx-4* and *xbx-4::gfp* sequences were expressed under the *xbx-4* upstream or *gpa-4Δ6* promoters. The *bbs-8*p::*dyf-18::gfp* (PSAB1121) and *bbs-8*p::*dyf-18 (KD)::gfp* (PSAB1124) constructs for overexpression have been described previously (11).

### Statistical analyses

Statistical analyses were performed using the SPSS 25 statistical analyses package (IBM). Phenotypic categories of cilia were defined as ordinal variables. The nonparametric Kruskal-Wallis test was used for data with non-normal distributions. When necessary, posthoc corrections for multiple comparisons were performed; the test used is indicated in each Figure Legend. Data are shown from at least 2-3 biologically independent experiments. Neither data acquisition nor analyses were blinded. Total numbers of analyzed cilia and significance values are indicated in each Figure Legend.

## Acknowledgements

We thank the *Caenorhabditis* Genetics Center and Shohei Mitani (National BioResource Project, Japan) for strains. We thank the Sengupta lab for advice and Hannah Lawson, Lauren Tereshko, Inna Nechipurenko and Oliver Blacque for comments on the manuscript. This work was funded in part by the NIH (R35 GM122463 – P.S.).

## Supplemental Figure Legends

**Figure S1.**
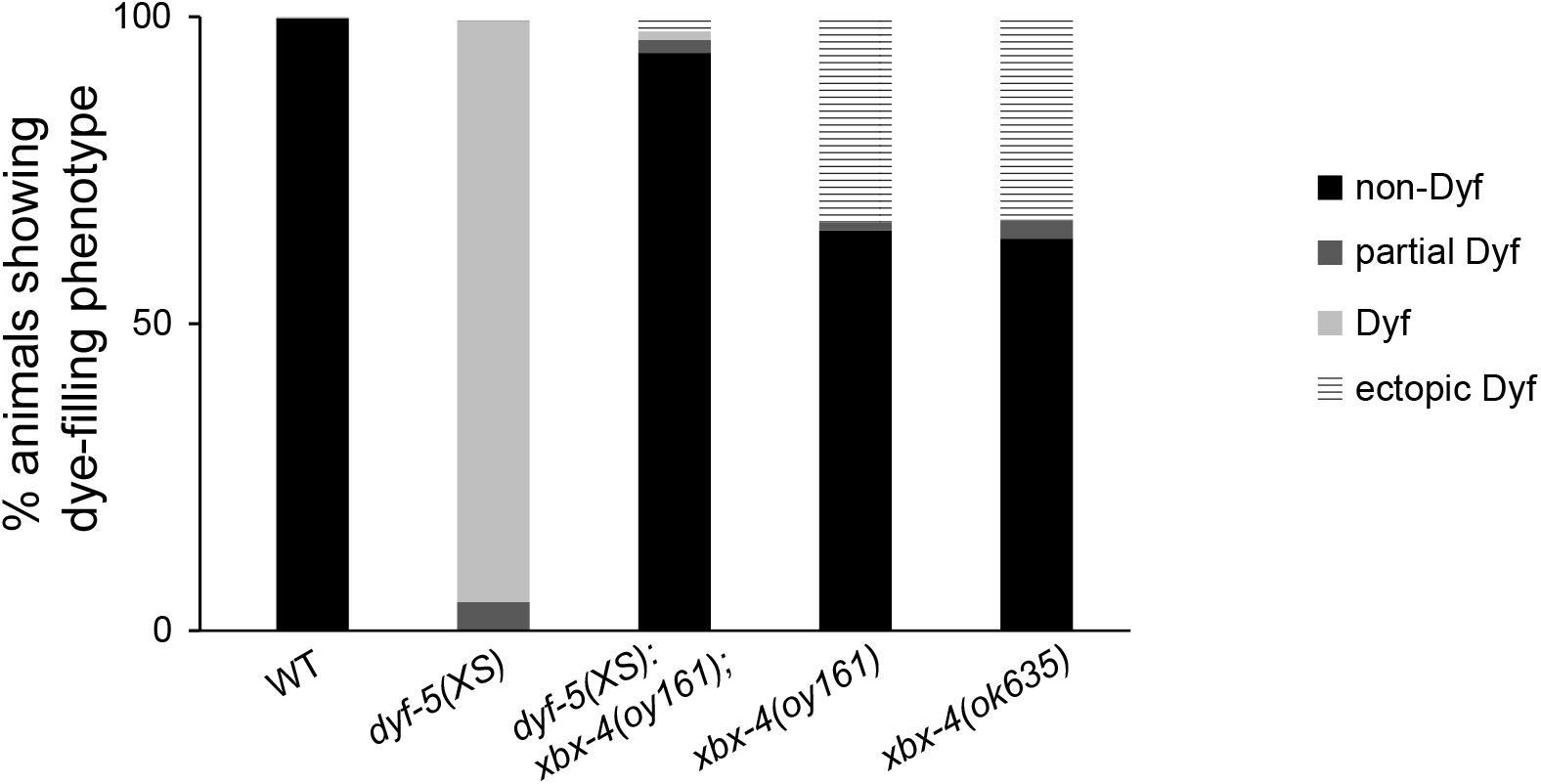
Mutations in *xbx-4* restore dye-filling to *dyf-5(XS)* mutants. Percentage of animals exhibiting dye-filling in six head amphid sensory neuron pairs in the indicated strains. nDyf: no dye-filling defect; partial Dyf: dye-filling observed in a subset of neurons; Dyf: no dye uptake; ectopic Dyf: dye uptake observed in additional head neurons (the dendritic bundle also appears unfasciculated in these animals). n≥300 animals each.

**Figure S2.**
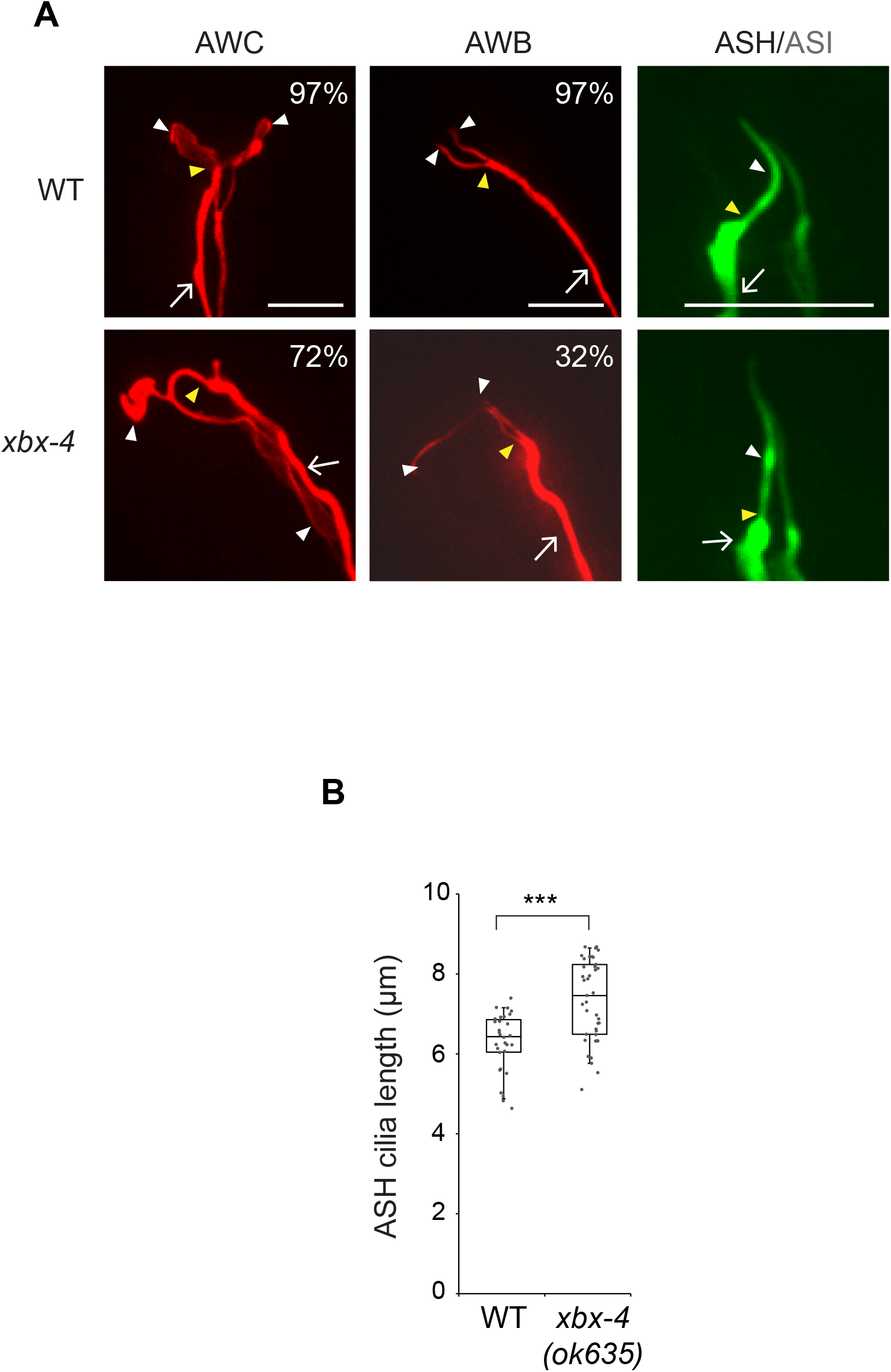
XBX-4 regulates length and morphology of multiple sensory cilia. **A)**Representative images of the cilia of the indicated sensory neurons in WT and *xbx-4(ok635)* mutants. AWC, AWB and ASH/ASI were visualized using *odr-1*p*::rfp, str-1*p*::mCherry* and *sra-6*p*::gfp*, respectively. Numbers at top right indicate the percentage of neurons showing the phenotype. Anterior at top left. n≥30 neurons each. **B)**Quantification of ASH cilium length. Each dot is the length of a single ASH cilium. Horizontal bars indicate 75^th^, 50^th^, and 25^th^ percentiles; error bars are 95^th^ and 5^th^ percentiles.

**Figure S3.**
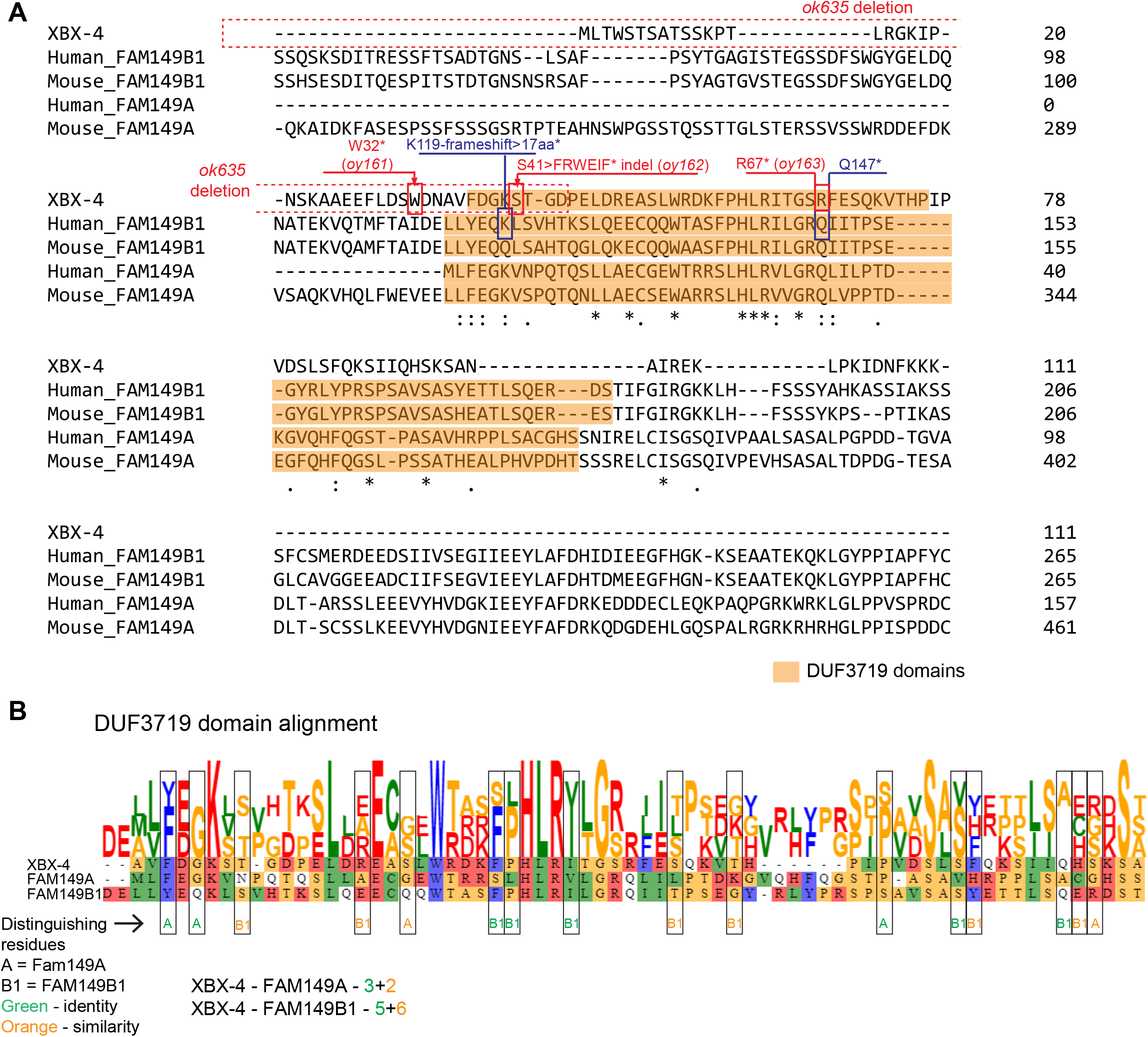
XBX-4 contains a conserved DUF3719 domain. **A)**Alignment of *C. elegans* XBX-4 and human and mouse FAM149A and FAM149B1 protein sequences. The DUF3719 domain in each protein as predicted by Interpro (https://www.ebi.ac.uk/interpro/) is shaded orange. The extent of the *ok635* deletion and locations and lesions of *xbx-4* mutations are indicated in red. Identified mutations in Joubert syndrome patients (4) are indicated in blue. **B)**Alignment of *C. elegans* and human DUF3719 domains (https://toolkit.tuebingen.mpg.de/tools/clustalo). To assess homology between XBX-4 and either of FAM149A or FAM149B1 DUF3719 domains, residues that are conserved between either XBX-4 and FAM149A, or XBX-4 and FAM149B1 (distinguishing residues) are indicated by boxes and color coded for identities or similarities.

**Figure S4.**
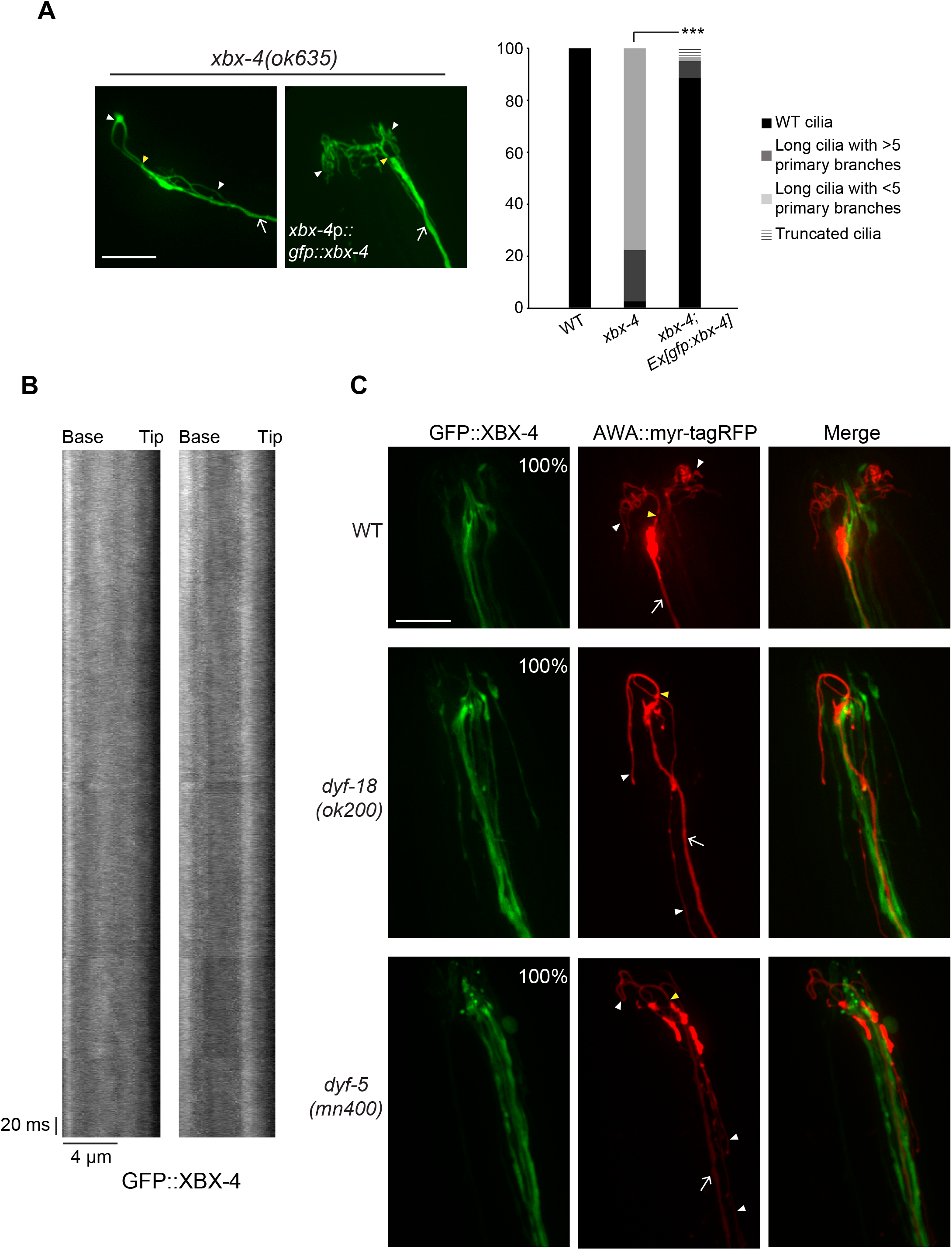
DYF-18 and DYF-5 do not regulate XBX-4 localization. **A)**GFP::XBX-4 rescues the AWA cilia morphology phenotype of *xbx-4(ok635)* mutants. Representative images and quantification of wild-type or *xbx-4(ok635)* mutants expressing a *xbx-4*p*:: gfp*::*xbx-4* transgene. AWA was visualized using *gpa-4Δ6*p*::myr-gfp.* Anterior at top left. ***: different between indicated at *P*<0.001 (Kruskal-Wallis test with Bonferroni post hoc corrections for multiple comparisons). n≥30 neurons each. **B)**Two representative kymographs of GFP::XBX-4 movement in channel cilia. **C)**Representative images of GFP::XBX-4 localization in the indicated genetic backgrounds. GFP::XBX-4 and myr-tagRFP were expressed under the *xbx-4* and *gpa-4Δ6* promoters, respectively. Numbers at top right indicate the percentage of neurons showing the phenotype. n≥20 neurons each. In all images, arrows: dendrite, yellow and white arrowheads: cilia base and cilia, respectively. Anterior at top left. Scale bar: 10 μm.

**Table S1.**
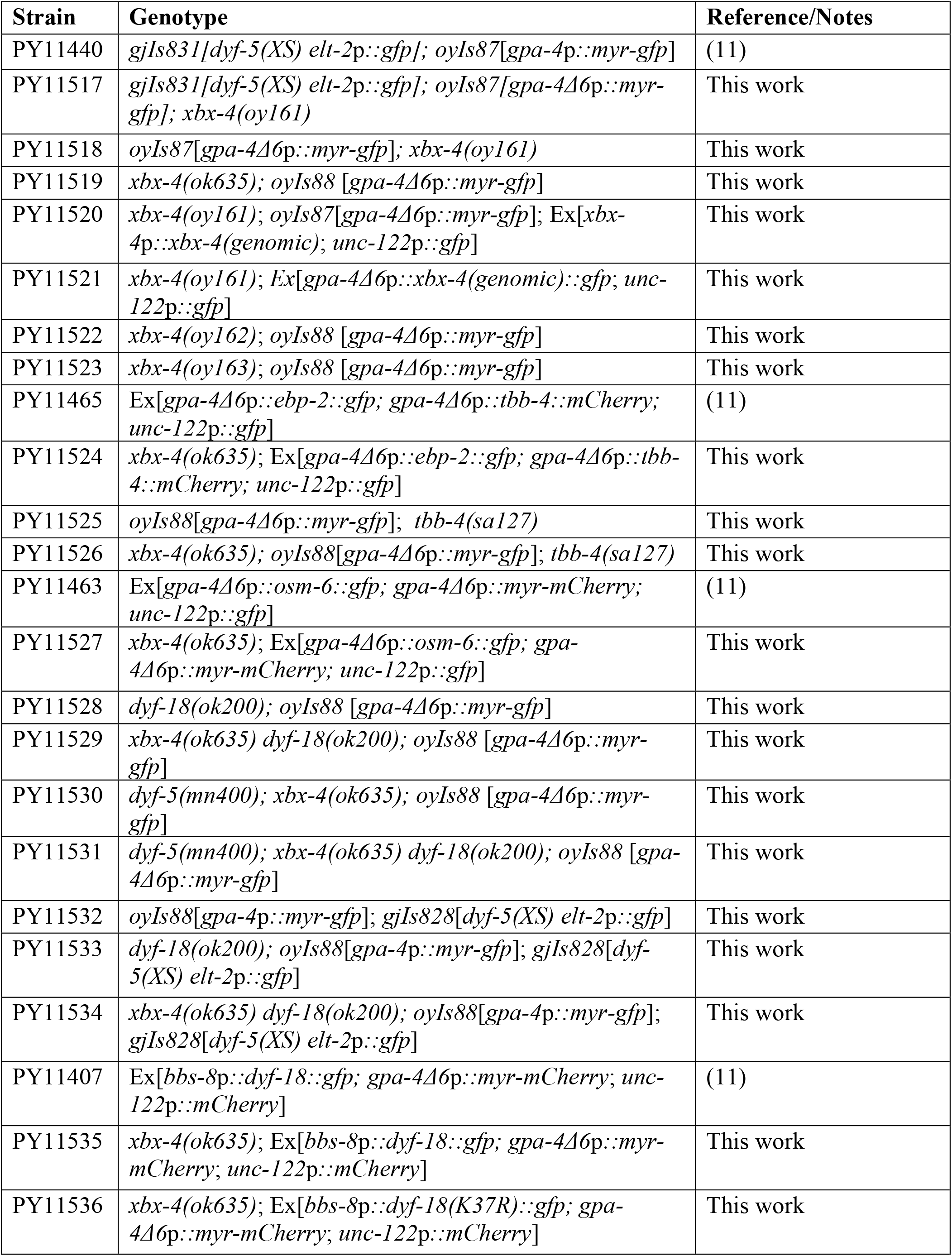
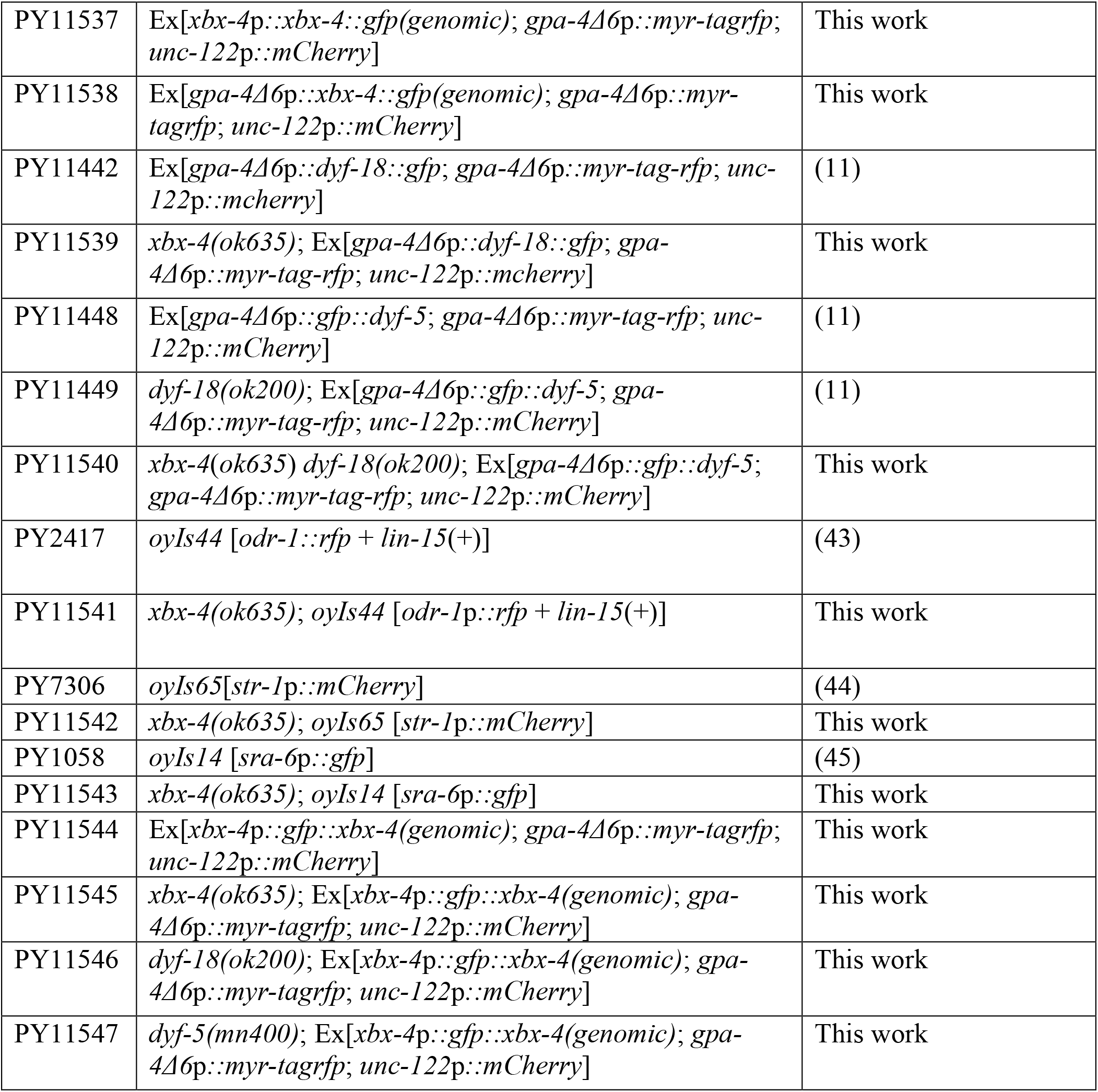
Strains used in this work.

**Table S2.** Plasmids used to generate strains in this work.

| **Plasmid identifier** | **Description** | **Source** |
| --- | --- | --- |
| PSAB1270 | *xbx-4p::xbx-4(genomic)* | This work |
| PSAB1271 | *gpa-4Δ6p::xbx-4(genomic)::gfp* | This work |
| PSAB1272 | *xbx-4p::xbx-4(genomic)::gfp* | This work |
| PSAB1133 | *gpa-4Δ6p::myr-tag-rfp* | (11) |
| PSAB1273 | *xbx-4p::gfp::xbx-4(genomic)* | This work |

## REFERENCES

1. Youn YH, Han YG. Primary cilia in brain development and diseases. Am J Pathol. 2018;188: 11–22.

2. Reiter JF, Leroux MR. Genes and molecular pathways underpinning ciliopathies. Nat Rev Mol Cell Biol. 2017;18: 533–547.

3. Benmerah A, Durand B, Giles RH, Harris T, Kohl L, Laclef C, et al. The more we know, the more we have to discover: an exciting future for understanding cilia and ciliopathies. Cilia. 2015;4: 5.

4. Shaheen R, Jiang N, Alzahrani F, Ewida N, Al-Sheddi T, Alobeid E, et al. Bi-allelic mutations in FAM149B1 cause abnormal primary cilium and a range of ciliopathy phenotypes in humans. Am J Hum Genet. 2019;104: 731–737.

5. Burghoorn J, Dekkers MP, Rademakers S, de Jong T, Willemsen R, Jansen G. Mutation of the MAP kinase DYF-5 affects docking and undocking of kinesin-2 motors and reduces their speed in the cilia of *Caenorhabditis elegans*. Proc Natl Acad Sci USA. 2007;104: 7157–7162.

6. Asleson CM, Lefebvre PA. Genetic analysis of flagellar length control in *Chlamydomonas reinhardtii*: a new long-flagella locus and extragenic suppressor mutations. Genetics. 1998;148: 693–702.

7. Berman SA, Wilson NF, Haas NA, Lefebvre PA. A novel MAP kinase regulates flagellar length in *Chlamydomonas*. Curr Biol. 2003;13: 1145–1149.

8. Tam LW, Wilson NF, Lefebvre PA. A CDK-related kinase regulates the length and assembly of flagella in *Chlamydomonas*. J Cell Biol. 2007;176: 819–829.

9. Moon H, Song J, Shin JO, Lee H, Kim HK, Eggenschwiller JT, et al. Intestinal cell kinase, a protein associated with endocrine-cerebro-osteodysplasia syndrome, is a key regulator of cilia length and Hedgehog signaling. Proc Natl Acad Sci USA. 2014;111: 8541–8546.

10. Broekhuis JR, Verhey KJ, Jansen G. Regulation of cilium length and intraflagellar transport by the RCK-kinases ICK and MOK in renal epithelial cells. PLoS One. 2014;9: e108470.

11. Maurya AK, Rogers T, Sengupta P. A CCRK and a MAK kinase modulate cilia branching and length via regulation of axonemal microtubule dynamics in *Caenorhabditis elegans*. Curr Biol. 2019;22: 1286–1300.

12. Herman RK, Hedgecock EM. Limitation of the size of the vulval primordium of *Caenorhabditis elegans* by *lin-15* expression in surrounding hypodermis. Nature. 1990;348: 169–171.

13. Starich TA, Herman RK, Kari CK, Yeh W-H, Schackwitz WS, Schuyler MW, et al. Mutations affecting the chemosensory neurons of *Caenorhabditis elegans*. Genetics. 1995;139: 171–188.

14. Perkins LA, Hedgecock EM, Thomson JN, Culotti JG. Mutant sensory cilia in the nematode *Caenorhabditis elegans*. Dev Biol. 1986;117: 456–487.

15. Doroquez DB, Berciu C, Anderson JR, Sengupta P, Nicastro D. A high-resolution morphological and ultrastructural map of anterior sensory cilia and glia in *C. elegans*. eLife. 2014;3: e01948.

16. Doitsidou M, Poole RJ, Sarin S, Bigelow H, Hobert O. *C. elegans* mutant identification with a one-step whole-genome-sequencing and SNP mapping strategy. PLoS ONE. 2010;5: e15435.

17. Efimenko E, Bubb K, Mak HY, Holzman T, Leroux MR, Ruvkun G, et al. Analysis of *xbx* genes in *C. elegans*. Development. 2005;132: 1923–1934.

18. Yi P, Xie C, Ou G. The kinases male germ cell-associated kinase and cell cycle-related kinase regulate kinesin-2 motility in *Caenorhabditis elegans* neuronal cilia. Traffic. 2018;19: 522–535.

19. Swoboda P, Adler HT, Thomas JH. The RFX-type transcription factor DAF-19 regulates sensory neuron cilium formation in *C. elegans*. Mol Cell. 2000;5: 411–421.

20. Hao L, Thein M, Brust-Mascher I, Civelekoglu-Scholey G, Lu Y, Acar S, et al. Intraflagellar transport delivers tubulin isotypes to sensory cilium middle and distal segments. Nat Cell Biol. 2011;13: 790–798.

21. Fu Z, Schroeder MJ, Shabanowitz J, Kaldis P, Togawa K, Rustgi AK, et al. Activation of a nuclear Cdc2-related kinase within a mitogen-activated protein kinase-like TDY motif by autophosphorylation and cyclin-dependent protein kinase-activating kinase. Mol Cell Biol. 2005;25: 6047–6064.

22. Fu Z, Larson KA, Chitta RK, Parker SA, Turk BE, Lawrence MW, et al. Identification of yin-yang regulators and a phosphorylation consensus for male germ cell-associated kinase (MAK)-related kinase. Mol Cell Biol. 2006;26: 8639–8654.

23. Tam LW, Dentler WL, Lefebvre PA. Defective flagellar assembly and length regulation in LF3 null mutants in *Chlamydomonas*. J Cell Biol. 2003;163: 597–607.

24. Nguyen RL, Tam LW, Lefebvre PA. The LF1 gene of *Chlamydomonas reinhardtii* encodes a novel protein required for flagellar length control. Genetics. 2005;169: 1415–1424.

25. Boldt K, van Reeuwijk J, Lu Q, Koutroumpas K, Nguyen TM, Texier Y, et al. An organelle-specific protein landscape identifies novel diseases and molecular mechanisms. Nat Commun. 2016;7: 11491.

26. Huttlin EL, Bruckner RJ, Paulo JA, Cannon JR, Ting L, Baltier K, et al. Architecture of the human interactome defines protein communities and disease networks. Nature. 2017;545: 505–509.

27. Mendes Maia T, Gogendeau D, Pennetier C, Janke C, Basto R. Bug22 influences cilium morphology and the post-translational modification of ciliary microtubules. Biology open. 2014;3: 138–151.

28. Yanagisawa HA, Mathis G, Oda T, Hirono M, Richey EA, Ishikawa H, et al. FAP20 is an inner junction protein of doublet microtubules essential for both the planar asymmetrical waveform and stability of flagella in *Chlamydomonas*. Mol Biol Cell. 2014;25: 1472–1483.

29. Laligne C, Klotz C, de Loubresse NG, Lemullois M, Hori M, Laurent FX, et al. Bug22p, a conserved centrosomal/ciliary protein also present in higher plants, is required for an effective ciliary stroke in *Paramecium*. Eukaryot Cell. 2010;9: 645–655.

30. Ko HW, Norman RX, Tran J, Fuller KP, Fukuda M, Eggenschwiler JT. Broad-minded links cell cycle-related kinase to cilia assembly and hedgehog signal transduction. Dev Cell. 2010;18: 237–247.

31. Snouffer A, Brown D, Lee H, Walsh J, Lupu F, Norman R, et al. Cell cycle-related kinase (CCRK) regulates ciliogenesis and Hedgehog signaling in mice. PLoS Genet. 2017;13: e1006912.

32. Hartwell LH, Hopfield JJ, Leibler S, Murray AW. From molecular to modular cell biology. Nature. 1999;402: C47–52.

33. Pereira-Leal JB, Levy ED, Teichmann SA. The origins and evolution of functional modules: lessons from protein complexes. Philos Trans R Soc Lond B Biol Sci. 2006;361: 507–517.

34. Cardenas-Rodriguez M, Badano JL. Ciliary biology: understanding the cellular and genetic basis of human ciliopathies. Am J Hum Med Genet. 2009;151C: 263–280.

35. Novarino G, Akizu N, Gleeson JG. Modeling human disease in humans: the ciliopathies. Cell. 2011;147: 70–79.

36. Park K, Li C, Tsiropoulou S, Goncalves J, Kondratev C, Pelletier L, et al. CDKL kinase regulates the length of the ciliary proximal segment. Curr Biol. 2021: S0960-9822(21)00441–3.

37. Lange KI, Tsiropoulou S, Kucharska K, Blacque OE. Interpreting the pathogenicity of Joubert Syndrome missense variants in *Caenorhabditis elegans*. Dis Models Mech. 2021;14: dmm046631.

38. Huang L, Szymanska K, Jensen VL, Janecke AR, Innes AM, Davis EE, et al. TMEM237 is mutated in individuals with a Joubert syndrome related disorder and expands the role of the TMEM family at the ciliary transition zone. Am J Hum Genet. 2011;89: 713–730.

39. Chen N, Mah A, Blacque OE, Chu J, Phgora K, Bakhoum MW, et al. Identification of ciliary and ciliopathy genes in *Caenorhabditis elegans* through comparative genomics. Genome Biol. 2006;7: R126.

40. Frokjaer-Jensen C, Davis MW, Sarov M, Taylor J, Flibotte S, LaBella M, et al. Random and targeted transgene insertion in *Caenorhabditis elegans* using a modified Mos1 transposon. Nat Methods. 2014;11: 529–534.

41. Minevich G, Park DS, Blankenberg D, Poole RJ, Hobert O. CloudMap: a cloud-based pipeline for analysis of mutant genome sequences. Genetics. 2012;192: 1249–1269.

42. Paix A, Folkmann A, Rasoloson D, Seydoux G. High efficiency, homology-directed genome editing in *Caenorhabditis elegans* using CRISPR-Cas9 ribonucleoprotein complexes. Genetics. 2015;201: 47–54.

43. Lanjuin A, VanHoven MK, Bargmann CI, Thompson JK, Sengupta P. *Otx/otd* homeobox genes specify distinct sensory neuron identities in *C. elegans*. Dev Cell. 2003;5: 621–633.

44. Nechipurenko IV, Olivier-Mason A, Kazatskaya A, Kennedy J, McLachlan IG, Heiman MG, et al. A conserved role for Girdin in basal body positioning and ciliogenesis. Dev Cell. 2016;38: 493–506.

45. Sarafi-Reinach TR, Melkman T, Hobert O, Sengupta P. The *lin-11* LIM homeobox gene specifies olfactory and chemosensory neuron fates in *C. elegans*. Development. 2001;128: 3269–3281.

